# Validation of TRIzol-Based Inactivation Protocol with Failure Scenario Testing for Bacterial Select Agent Surrogates

**DOI:** 10.64898/2026.02.22.707045

**Authors:** Umar Raza Shahid, Paul Augustus Lueth, Bryan H. Bellaire

## Abstract

Validated inactivation procedures are required for the safe handling and downstream analysis of highly pathogenic organisms, particularly those categorized as biological select agents and toxins (BSATs). TRIzol™-based extraction methods are widely used for nucleic acid and protein isolation, yet their reliability for bacterial inactivation has not been comprehensively evaluated. In this study, we assessed TRIzol-based extraction methods for sample quality and inactivation reliability across a series of mock failure scenarios using five attenuated bacterial isolates: *Francisella tularensis holarctica* LVS, *Bacillus anthracis* Sterne, *Yersinia enterocolitica, Mycobacterium marinum*, and *Burkholderia cepacia*. Dilution of TRIzol to induce incomplete cell lysis for the initial extraction step, including 0% TRIzol, consistently inactivated all surrogate organisms, suggesting that downstream precipitation and sample washing reagents, including isopropanol and 70% ethanol, were sufficient to inactivate organisms in the absence of TRIzol. Several protocol failure scenarios were then evaluated to simulate human error by omitting extraction, precipitation, and washing steps individually or in combination for the most resistant organism, *B. anthracis* Sterne strain. Failure-scenario testing demonstrated that reliable inactivation of *B. anthracis* required strict adherence to the complete protocol due to the spore-forming ability of *B. anthracis*. Collectively, this work provides a reference with experimental evidence supporting the use of TRIzol™-based extraction as a bacterial inactivation strategy for a wide range of bacterial pathogens.

## Introduction

The term “Select Agents” refers to specific biological agents and toxins that could harm public, animal, or plant health. Under the regulatory framework standardization established by the United States Department of Health and Human Services (HHS) and the United States Department of Agriculture (USDA), codified in 42 CFR Part 73, 7 CFR Part 331, and 9 CFR Part 121, laboratories working with select agents must implement validated inactivation procedures that demonstrate the non-viability of treated materials through comprehensive viability testing protocols[3] [1], [2].

Nucleic acid extraction represents a critical component of select agent research, enabling molecular characterization while requiring complete inactivation of the source organism. The Federal Select Agent Program (FSAP) recognizes that, while 100% inactivation is neither mathematically nor practically achievable, the risk of viable agents in processed materials must be reduced to the lowest possible level, given the intended use and the potential consequences of inactivation failures [3]. This imperative has driven the development and validation of standardized nucleic acid extraction protocols, including phenol-chloroform-based methods such as TRIzol, which combine chemical lysis with purification steps to achieve dual objectives of RNA, DNA and protein isolation and pathogen inactivation [4], [5]. While multiple studies have described TRIzol™-based inactivation methods for BSAT viruses, there is a notable lack of validated protocols for bacterial pathogens [5], [6], [7].

The complexity of validating inactivation protocols across diverse microbial agents necessitates the use of appropriate surrogate organisms [3] that represent the range of resistance characteristics encountered in actual select agents. Bacterial surrogates, including *Francisella tularensis* subsp. *holartica* LVS, *Bacillus anthracis, Yersinia enterocolitica, Mycobacterium marinum*, and *Burkholderia cepacia* provide a comprehensive model system representing gram-negative bacteria, spore-forming organisms, and acid-fast bacteria with varying degrees of environmental resistance and cell wall complexity.

### Study Rationale and Objectives

This study addresses the critical need for comprehensive validation of TRIzol-based inactivation protocols specifically designed to withstand common failure scenarios encountered in select agent laboratories. Working in partnership with regulatory agencies to establish standardized protocols for broader adoption, this research systematically challenges established procedures with controlled failure conditions.

The primary objectives of this research are to: (1) establish a validated TRIzol-based inactivation protocol for broth culture samples using a panel of surrogate organisms (*F. tularensis, B. anthracis, Y. enterocolitica, M. marinum*, and *B. cepacia*) that represent the diversity of select agents, (2) systematically test the protocol against various failure scenarios including procedural deviations, reagent omissions, and potential matrix interferences, and (3) quantify the safety margins inherent in the optimized protocol across different bacterial morphotypes and resistance characteristics.

## Materials and Methods

### Bacterial Strains and Culture Conditions

In accordance with Federal Select Agent Program (FSAP) guidelines, surrogate BSL-2 strains were used as safe alternatives to their BSL-3 pathogenic counterparts. The following surrogate laboratory-derived organisms were employed: (1) *Yersinia enterocolitica* for *Y. pestis*, (2) *Bacillus anthracis* Sterne strain for virulent *B. anthracis*, (3) *Francisella tularensis* subsp. *holarctica* LVS (NR-646) for *F. tularensis* subsp. *tularensis*, (4) *Mycobacterium marinum* for *M. tuberculosis*, (5) *Burkholderia cepacia* (ATCC NR-707) for *Burkholderia pseudomallei*. All five surrogate bacterial strains were successfully cultured under their respective optimal conditions. Initial cultures were grown on agar media started from frozen (-80ºC). A single colony was transferred by inoculation loop into 10 mL of broth medium and cultured as indicated for each strain (Table 1). Initial bacterial concentrations ranged from 5.3 × 10^6^ to 1.6 × 10^9^ CFU/mL across all strains.

**Table 1.** Bacterial strains, respective culture media and bacterial densities for TRIzol extraction.

| Surrogate/strain | Temp | Broth | Agar | (CFU/mL) |
| --- | --- | --- | --- | --- |
| *Yersinia enterocolitis* | 26 ° C | Tryptic Soy Broth | Hb-Columbia Agar | $1.6 \times 10^9$ |
| *Bacillus anthracis* Sterne strain | 37 ° C | Tryptic Soy Broth | Hb-Columbia Agar | $1.2 \times 10^8$ |
| *Francisella tularensis* subsp. *holarctica* LVS | 37 ° C | Tryptic Soy Broth | Hb-Columbia Agar | $3 \times 10^8$ |
| *Mycobacterium marinum* | 30 ° C | 7H9 Broth | Hb-Columbia Agar | $5.3 \times 10^6$ |
| *Burkholderia cepacia* | 37 ° C | Mueller Hinton Broth | 5% Sheep Blood Agar | $1.5 \times 10^8$ |

### Inactivation Test by Recovery of Viable Organisms

Confirmation of inactivation from the final DNA, RNA, and protein extractions by transferring 50 µL of the final product into 10mL fresh broth and incubated for seven days under appropriate conditions. Subsequently, 1 mL of the seven-day culture was plated onto agar media and incubated for an additional seven days to assess bacterial viability. Samples devoid of colonies on agar plates following incubation were recorded as inactivated. Culture media and incubation conditions for each strain are detailed in Table 1.

### Standard TRIzol™-Based Nucleic Acid and Protein Extraction

The objective of this study was to evaluate the efficiency of inactivation of Select Agents using a standardized protocol for biosafe RNA, DNA, and protein extraction. from select agents to allow for transfer outside of processing outside FSAP-controlled facilities. The extraction methodology employed TRIzol™ Reagent (Thermo Fisher Scientific) according to the manufacturer’s protocol, with modifications as described below. A schematic overview of the process is shown in Figure 1.

**Figure 1.**
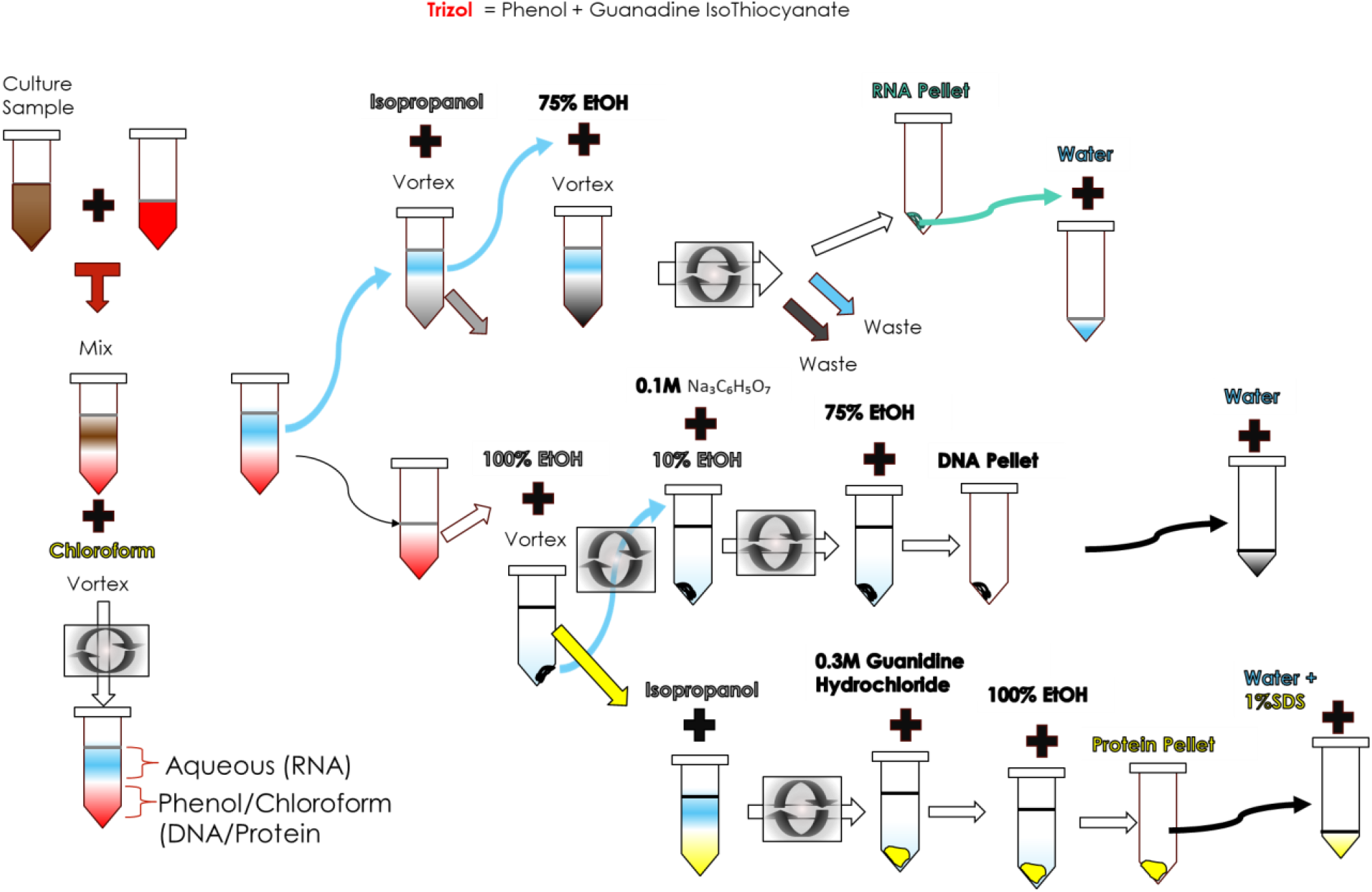
Overview of TRIzol based extraction and chemicals used.

Samples were prepared by transferring 500 µL of bacterial culture to a 2 mL microcentrifuge tube and centrifuging at 10,000 × g for 10 minutes. The supernatant was discarded, and the bacterial pellet was resuspended in 500 µL of TRIzol™ reagent. Samples were incubated at room temperature for 5 minutes to ensure complete cell lysis and nucleic acid dissociation. Aqueous and organic phases were separated by adding 100 µL of chloroform, vortexed, incubated at room temperature for 2-3 minutes, then centrifuged at 12,000 × g for 15 minutes at 4°C. Phase separation into three distinct layers: a colorless upper aqueous phase containing RNA, the interphase layer (containing DNA), and a lower phenol-chloroform phase (containing proteins). The upper aqueous phase containing RNA was carefully transferred to a new microcentrifuge tube with 250 µL of isopropanol for RNA precipitation facilitated by centrifugation. The RNA pellet was washed with 500 µL of 75% ethanol, centrifuged again, and air-dried in a biosafety cabinet for 5-10 minutes. The final RNA pellet was resuspended in 50 µL of RNase-free water and stored at -80°C. DNA was isolated from the interphase and phenol-chloroform phase by adding 150 µL of 100% ethanol, incubating for 2-3 minutes at room temperature. The mixture was centrifuged to pellet the DNA away from the proteins in the supernatant, for recovery as described below. The tube containing the DNA pellet was washed sequentially with 500 µL of 0.1 M sodium citrate in 10% ethanol for 30 minutes, followed by 500 µL of 75% ethanol for 10 minutes. After final centrifugation, the DNA pellet was air-dried and resuspended in 50 µL of 8 mM NaOH. Supernatants from the above step were precipitated by incubation with 750 µL of isopropanol for 10 minutes at room temperature, followed by centrifugation. The protein pellet was washed twice with 300 µL of 0.3 M guanidine hydrochloride in 95% ethanol for 20 minutes each, followed by a final wash with 1 mL of 100% ethanol. After air-drying, the pellet was resuspended in 50 µL of 1% SDS. Nucleic acid concentrations and purity were determined using a NanoDrop 2000 spectrophotometer (Thermo Fisher Scientific), with A_260_/A_280_ ratios recorded to assess purity. DNA, RNA, and protein purity were quantified by UV absorbance. All extractions were performed in triplicate.

### Failure by dilution of TRIzol™

To establish minimum effective concentrations for bacterial inactivation, five different concentrations of TRIzol™ Reagent were evaluated. All dilutions were prepared using sterile double-distilled water (ddH_2_O) as follows: Full strength: 100% TRIzol™ Reagent, 75% TRIzol™ Reagent, 1/2 strength: 50% TRIzol™ Reagent, 1/4 strength: 25% TRIzol™ Reagent and Negative control: 0% TRIzol™ Reagent or water only. Remaining steps were fodescribed in initial protocol and in triplicate against all bacterial strains.

### Failure by step-wise omission for *B. anthracis* Sterne strain

*Bacillus anthracis* Sterne strain inactivation was further evaluated by failure to conduct individual step scenario was tested only with as prior results demonstrated that the omission of TRIzol caused the only failure in the test indicating that the downstream processing steps were insufficient. To identify the steps that contributed the greatest inactivation potential, we designed a failure test based on progressive step omission. Beginning with TRIzol omission by adding bacteria directly to chlororom and completing the RNA isolation. The next sample represented omission of the first 2 steps by directly adding bacteria to isopropanol and compleing RNA isolation. This format was carried out for the remaining steps, in total 4 conditions were evaluated for *B. anthracis* Sterne strain inactivation (Table 3).

### Inactivation by Treatment with Single Reagent

To assess the individual antimicrobial activity of each reagent used in the TRIzol™ protocol, single-reagent treatments were evaluated for *B. anthracis* and *B. abortus* Strain-19. Bacterial samples of 0.5 ml were mixed with indicated amount and time for each reagent tested. (TRIzol™ Reagent, Chloroform, Isopropanol or water only). Each treatment was applied using the same volumes, incubation times, and conditions as specified in the complete protocol by the manufacturer to maintain consistency and enable direct comparison of antimicrobial effectiveness. At the end of incubation time, the bacterial suspension was centrifuged, and the pellet was transferred to a 10 mL culture tube for 7-day incubation. Then, 1 mL was spread on an agar plate and further incubated for 7 days. All samples were processed in triplicate. Positive and negative growth control for bacterial species were also performed.

## Results

### Standard TRIzol™-Based Nucleic Acid and Protein Extraction

Complete TRIzol™ extraction of RNA, DNA and protein protocol was evaluated for quality of sample isolation and efficiency of bacterial inactivation for each sample type for each bacterial species. For all seven bacterial species, 100% bacterial inactivation was achieved for all Trizol samples with the exception of the DNA sample recovered from *B. anthracis* Sterne strain. Inactivation test method reflects the absence of viable bacteria recovered after 7-day incubation in broth or subsequent plating on agar media. The 0% TRIzol samples also showed full inactivation of RNA, DNA, and protein extraction, with the exception that two out of three (2/3) *B. anthracis* DNA samples failed inactivation and showed growth on Hb-Columbia agar. Later, the CFU/ml was quantified from 7-day broth (Figure 2). In the absence of TRIzol, the remaining precipitation and washing solutions of isoproponal, alcohol and chloroform that are known antimicrobial and disinfection solutions, were sufficient in inactivating bacterial samples for all except *B. anthracis* Sterne DNA sample. Table 2.

**Table 2.** Assessment of nucleic acid and protein recovery following TRIzol treatment at varying concentrations. Five bacterial species (*Yersinia enterocolitica, Bacillus anthracis* Sterne strain, *Francisella tularensis* subsp. *holarctica* LVS, *Mycobacterium marinum*, and *Burkholderia cepacia*) were exposed to 100%, 75%, 50%, 25%, and 0% TRIzol. Successful recovery of DNA (D), RNA (R), and protein (P) was evaluated across three independent replicates per condition and reported as the number of successful recoveries out of three trials (e.g., 3/3). Results demonstrate consistent recovery across treatments, with B. anthracis Sterne at variation.

|  | 100% TRIzol |  |  | 75% TRIzol |  |  | 50% TRIzol |  |  | 25% TRIzol |  |  | 0% TRIzol |  |  |
| --- | --- | --- | --- | --- | --- | --- | --- | --- | --- | --- | --- | --- | --- | --- | --- |
|  | D | R | P | D | R | P | D | R | P | D | R | P | D | R | P |
| *Yersinia enterocolitica* | 3/3 | 3/3 | 3/3 | 3/3 | 3/3 | 3/3 | 3/3 | 3/3 | 3/3 | 3/3 | 3/3 | 3/3 | 3/3 | 3/3 | 3/3 |
| *Bacillus anthracis* Sterne strain | 3/3 | 3/3 | 3/3 | 3/3 | 3/3 | 3/3 | 3/3 | 3/3 | 3/3 | 3/3 | 3/3 | 3/3 | 2/3 | 3/3 | 3/3 |
| *Francisella tularensis* subsp. *holarctica* LVS | 3/3 | 3/3 | 3/3 | 3/3 | 3/3 | 3/3 | 3/3 | 3/3 | 3/3 | 3/3 | 3/3 | 3/3 | 3/3 | 3/3 | 3/3 |
| *Mycobacterium marinum* | 3/3 | 3/3 | 3/3 | 3/3 | 3/3 | 3/3 | 3/3 | 3/3 | 3/3 | 3/3 | 3/3 | 3/3 | 3/3 | 3/3 | 3/3 |
| *Burkholderia cepacia* | 3/3 | 3/3 | 3/3 | 3/3 | 3/3 | 3/3 | 3/3 | 3/3 | 3/3 | 3/3 | 3/3 | 3/3 | 3/3 | 3/3 | 3/3 |

**Table 3.** Inactivation results following stepwise failure scenarios. All scenarios omitting Trizol failed to inactivate any of the triplicate samples tested for each condition.

| Condition | Steps included in failure scenario |  |  |  |  | Inactivated % |
| --- | --- | --- | --- | --- | --- | --- |
| All Steps | TRizol | Chloroform | Isopropanol | 75% alcohol | Water | 100% |
| Four |  | Chloroform | Isopropanol | 75% alcohol | Water | 0% |
| Three |  |  | Isopropanol | 75% Alcohol | Water | 0% |
| Two |  |  |  | 75% Alcohol | Water | 0% |
| One |  |  |  |  | Water | 0% |

**Table 4.** Individual Reagent Inactivation Efficacy Results: Trizol reagent consistently achieved complete inactivation of B. anthracis Sterne Strain. No other individual regent achieved inactivation.

| Chemical | Volume | Time of incubation | Inactivation %<br>*B. anthracis* |
| --- | --- | --- | --- |
| TRizol | 500 µL | 5 min | 100% |
| Chloroform | 100 µL | 2-3 min | 0% |
| Isopropanol | 250 µL | 10 min | 0% |
| 75% ethanol | 500 µL | 5 min | 0% |
| Water | 50 µL | ---- | 0% |

The impact on quality of the DNA, RNA and protein samples harvested correlated with percentage of Trizol in first step. DNA extraction resulted in concentrations from 3.213 to 451.0 ng/µL across strains. ¾ strength TRIzol yielded the highest average DNA concentration, followed by ¼ strength and full strength. The A_260_/A_280_ ratios indicated varying purity levels, suggesting Purity metrics were affected by phenol carryover inherent to TRIzol-based extraction. Higher values were noted with dilutions below ½ strength TRIzol. RNA extraction across all strains, with concentrations from 581 to 0 ng/μL. Full strength TRIzol also produced the highest mean RNA concentration, followed by ¾ strength. The A_260_/A_280_ ratios indicated acceptable RNA purity for all strains except *B. cepacia* no TRIzol samples. Protein concentrations varied from 0.14 to 4.66 mg/mL. The observed overall trend was that higher amounts and quality recovered corresponded to higher amounts of Trizol likely due to more complete cell lysis (Supplemental 1).

### Sequential Reagent Omission Analysis and Individual Reagent Inactivation Efficacy

The ability of 0% Trizol treated samples to inactivate samples demonstrated that the remaining solutions used for precipitation and purification were sufficient at inactivating any remaining bacteria. To understand the added safety margin that is afforded by these steps, we evaluated the tolerance of omitting steps and assessing bacterial viability (Table 3). This scenario was evaluated the RNA isolation pathway using the *Bacillus anthracis* Sterne strain. Sequential omission of reagents demonstrated that complete adherence to the TRIzol™ protocol resulted in 100% bacterial inactivation. In contrast, omission of TRIzol™ and chloroform from the protocol resulted in inactivation failure demonstrating that TRIzol™ and chloroform is necessary for bacterial inactivation. A similar finding was observed when individual treatments were examined where only TRIzol was capable of inactivating *B. anthracis* samples (Table 3).

## Discussion

This study successfully demonstrates the efficacy of TRIzol-based inactivation protocols for diverse bacterial surrogates representing major select agent categories. The achievement of 100% inactivation for the tested strains validates the TRIzol protocol suitability for regulatory compliance under FSAP guidelines [3]. The single exception, *B. anthracis*, which required adherence to the complete protocol, underscores the critical importance of standardized procedures and highlights the protocol built-in safety margins when properly executed. These findings are in agreement with another research concerning the differential inactivation of *Bacillus anthracis* in comparison to other microorganisms during the sample preparation phase for Matrix-Assisted Laser Desorption/Ionization Time-of-Flight Mass Spectrometry (MALDI-TOF MS) [8], [9] and monochloramine inactivation in drinking water [10]. There is limited information available regarding effective chemical inactivation strategies for this organism [11].

The systematic evaluation of the individual TRIzol method of RNA extraction for *B. anthracis* components provides critical insights into the mechanistic basis of inactivation efficacy. 100% inactivation rate of TRIzol only confirms its role as the primary antimicrobial agent, consistent with its phenol-guanidinium thiocyanate composition that disrupts cellular membranes and denatures proteins.

The protocol demonstrated tolerance to TRIzol dilution but not to reagent omission, indicating that robustness is conditional upon inclusion of all critical reagents, with maintained inactivation efficacy across TRIzol concentrations, method modifications, and procedural deviations. This robustness addresses a critical regulatory concern identified in FSAP guidance [3], where human error represents a significant contributor to inactivation failures. The protocol’s tolerance for minor procedural variations while maintaining complete inactivation provides essential safety margins for routine laboratory implementation. However, the requirement for complete protocol adherence with *B. anthracis* serves as an important reminder that certain organisms may represent edge cases requiring strict procedural compliance. This finding supports FSAP’s emphasis on comprehensive validation across diverse microbial types and reinforces the importance of process controls and verification procedures for challenging organisms [3].

The successful recovery of DNA, RNA, and protein across all tested organisms confirms the protocol’s compatibility with diverse downstream analytical applications. The maintenance of nucleic acid and protein integrity while achieving complete pathogen inactivation addresses a fundamental challenge in select agent research, where analytical requirements must be balanced against safety imperatives. The protocol demonstrated compatibility with molecular techniques enables comprehensive characterization studies while maintaining regulatory compliance.

While this study focused on broth culture samples, the mechanistic basis of TRIzol inactivation suggests broad applicability across diverse sample types commonly encountered in select agent work. The protocol effectiveness against gram-negative bacteria, spore-forming organisms, and acid-fast bacteria indicates potential compatibility with tissue samples, and environmental specimens which should be tested separately. The established protocol framework can serve as a foundation for extending validation to additional sample types, with organism-specific optimization guided by the analysis framework developed in this study.

The comprehensive validation data generated in this study provides a strong foundation for regulatory submission and potential standardization across select agent laboratories. The systematic evaluation of failure scenarios, individual reagent contributions, and organism-specific effects addresses key concerns identified in FSAP guidance documents [3]. The protocol demonstrated remarkable robustness and broad organism compatibility suggest potential for adoption as a standardized method across the select agent community. The detailed characterization of critical control points and safety margins provides essential information for training programs and quality assurance procedures. The integration of this protocol into existing laboratory workflows should enhance both safety and analytical capabilities while maintaining regulatory compliance.

## Conclusions

This study establishes a validated TRIzol-based inactivation protocol that successfully balances pathogen inactivation requirements with analytical sample integrity across diverse bacterial surrogates. The TRIzol protocol demonstrated robustness under various failure scenarios, combined with downstream analytical compatibility, supports its adoption for routine select agent laboratory operations. The comprehensive characterization of individual reagent contributions and organism-specific effects provides essential guidance for protocol optimization and quality control procedures. The analysis framework developed in this study offers a quantitative approach to protocol validation that can be extended to additional organisms and sample types. The identification of critical control points and safety margins provides important insights for training programs and regulatory compliance procedures.

The successful validation of this protocol addresses critical needs identified in FSAP guidance while providing practical solutions for laboratory operations. The demonstrated compatibility with diverse analytical applications, combined with robust inactivation efficacy, supports the protocol’s potential for standardization across the select agent research community.

## Acknowledgments

[To be added as appropriate]

## Supplementary Materials

1. Excel file of RNA DNA and protein yields.

## Supplemental

**Supplemental Table 1:** RNA, DNA and Protein absorbance readings for quantity and quality for each replicate of samples corresponding to variable TRIzol testing.

| RNA |  | Smpl | Francisella tularensis<br>subsp. holarctica LVS |  |  | Bacillus anthracis<br>Sterne strain |  |  | Yersinia<br>enterocolitica |  |  | Mycobacterium<br>marinum |  |  | Burkholderia cepacia |  |  |
| --- | --- | --- | --- | --- | --- | --- | --- | --- | --- | --- | --- | --- | --- | --- | --- | --- | --- |
|  |  |  | ng/ul | 260/<br>280 | 260/<br>230 | ng/ul | 260/<br>280 | 260/<br>230 | ng/ul | 260/<br>280 | 260/<br>230 | ng/ul | 260/<br>280 | 260/<br>230 | ng/ul | 260/2<br>80 | 260/<br>230 |
| 100%<br>TRIZol |  | 1.1 | 95.55 | 1.75 | 1.52 | 55.29 | 1.69 | 1.69 | 655.86 | 1.67 | 0.5 | 530.37 | 2.23 | 0.16 | 141.6 | 0.72 | NA |
|  |  | 1.2 | 76.57 | 1.84 | 1.64 | 200.2 | 1.83 | 1.83 | 531.42 | 1.68 | 0.57 | 471.15 | 1.75 | 0.24 | 158.6 | 0.7 | NA |
|  |  | 1.3 | 177.24 | 1.82 | 0.49 | 353.84 | 2.04 | 2.04 | 558.12 | 1.74 | 0.38 | 588.03 | 1.72 | 0.46 | 119.9 | 0.63 | NA |
| 75%<br>Strengt<br>h<br>TRIZol |  | 2.1 | 51.27 | 1.78 | 0.87 | 485.72 | 1.55 | 1.55 | 366.68 | 1.93 | 0.75 | 31.99 | 1.81 | 0.12 | 154.6 | 0.63 | NA |
|  |  | 2.2 | 50.34 | 1.77 | 0.96 | 660.58 | 1.42 | 1.42 | 511.73 | 1.62 | 0.46 | 713.62 | 1.75 | 0.32 | 108.1 | 0.77 | NA |
|  |  | 2.3 | 50.21 | 1.77 | 0.72 | 601.49 | 1.67 | 1.67 | 425.54 | 2.04 | 0.86 | 233.28 | 1.91 | 0.44 | 171.6 | 0.72 | NA |
| 50%<br>Strengt<br>h<br>TRIZol |  | 3.1 | 13.85 | 1.77 | 0.24 | 282.04 | 1.93 | 1.93 | 587.71 | 1.63 | 0.42 | 96.46 | 1.74 | 0.2 | 273.6 | 1.01 | NA |
|  |  | 3.2 | 64.08 | 1.7 | 0.51 | 100.46 | 1.75 | 1.75 | 311.02 | 1.82 | 0.66 | 958.13 | 1.73 | 0.4 | 133.3 | 0.6 | NA |
|  |  | 3.3 | 72.96 | 1.6 | 0.4 | 99.47 | 1.73 | 1.73 | 357.83 | 1.87 | 0.7 | 159.45 | 1.8 | 0.33 | 127.4 | 0.63 | NA |
| 25%<br>Strengt<br>h<br>TRIZol |  | 4.1 | 52.49 | 1.73 | 0.96 | 85.71 | 1.72 | 1.72 | 176.07 | 1.66 | 0.61 | 245.21 | 1.87 | 0.46 | 223 | 0.89 | NA |
|  |  | 4.2 | 22.85 | 1.86 | 1.66 | 164.87 | 1.78 | 1.78 | 111.29 | 1.43 | 0.68 | 88.45 | 1.71 | 0.6 | 149.1 | 0.64 | NA |
|  |  | 4.3 | 45.33 | 1.71 | 0.95 | 38.53 | 1.71 | 1.71 | 182.57 | 1.68 | 0.57 | 61.55 | 1.68 | 0.16 | 198.3 | 0.77 | NA |
| 0%<br>Water<br>only |  | 5.1 | -2.12 | 0.77 | 1.18 | 3.64 | 1.69 | 1.69 | 142.76 | 1.17 | 0.79 | 133.61 | 1.22 | 0.88 | 109.3 | 1.46 | NA |
|  |  | 5.2 | -2.66 | 1.81 | 1.33 | 3.18 | 1.83 | 1.83 | 145.68 | 1.17 | 0.79 | 9.21 | 1.8 | 0.68 | 63 | 1.67 | NA |
|  |  | 5.3 | -2.02 | 1.32 | 2.27 | 1.72 | 29.9 | 29.9 | 199.37 | 1.26 | 0.87 | 15.95 | 1.6 | 0.9 | 110.3 | 41.58 | NA |

| DNA |  | Smpl | Francisella tularensis<br>subsp. holarctica LVS |  |  | Bacillus anthracis<br>Sterne strain |  |  | Yersinia<br>enterocolitica |  |  | Mycobacterium<br>marinum |  |  | Burkholderia cepacia |  |  |
| --- | --- | --- | --- | --- | --- | --- | --- | --- | --- | --- | --- | --- | --- | --- | --- | --- | --- |
|  |  |  | Con.<br>ng/ul | 260/<br>280 | 260/<br>230 | Con.<br>ng/ul | 260/<br>280 | 260/<br>230 | Con.<br>ng/ul | 260/<br>280 | 260/<br>230 | Con.<br>ng/ul | 260/2<br>80 | 260/<br>230 | Con.<br>ng/ul | 260/2<br>80 | 260/<br>230 |
| 100%<br>TRIZol |  | 1.1 | 73.73 | 0.84 | 0.36 | 6.34 | 0.23 | 0.06 | 179.42 | 1.24 | 0.74 | 180.07 | 0.78 | 0.34 | 141.6 | 0.72 | NA |
|  |  | 1.2 | 75.54 | 0.86 | 0.38 | 341.77 | 1.51 | 0.98 | 172.88 | 0.93 | 0.38 | 268.73 | 0.89 | 0.45 | 158.6 | 0.7 | NA |
|  |  | 1.3 | 104.8 | 0.98 | 0.45 | 684 | 1.4 | 1.02 | 103.26 | 0.65 | 0.25 | 226.86 | 1.49 | 0.45 | 119.9 | 0.63 | NA |
| 75%<br>Strength<br>TRIZol |  | 2.1 | 410.39 | 0.92 | 0.63 | 323.04 | 1.27 | 0.73 | 53.03 | 0.52 | 0.16 | 204.36 | 0.76 | 0.35 | 154.6 | 0.63 | NA |
|  |  | 2.2 | 61.2 | 0.63 | 0.24 | 749.75 | 1.17 | 0.69 | 872.65 | 1.42 | 0.8 | 130.92 | 0.59 | 0.24 | 108.1 | 0.77 | NA |
|  |  | 2.3 | 35.97 | 0.57 | 0.15 | 280.06 | 1.27 | 0.7 | 173.38 | 0.76 | 0.34 | 184.29 | 0.68 | 0.3 | 171.6 | 0.72 | NA |
| 50%<br>Strength<br>TRIZol |  | 3.1 | 47.1 | 0.57 | 0.22 | 339.11 | 1.16 | 0.63 | 208.21 | 0.96 | 0.4 | 95.94 | 0.52 | 0.19 | 273.6 | 1.01 | NA |
|  |  | 3.2 | 47.47 | 0.58 | 0.21 | 280.53 | 1.42 | 0.9 | 263.7 | 1.02 | 0.54 | 73.74 | 0.78 | 0.29 | 133.3 | 0.6 | NA |
|  |  | 3.3 | 166.29 | 1.43 | 0.83 | 425.15 | 1.29 | 0.81 | 61.96 | 0.48 | 0.16 | 125.9 | 4.507 | 0.56 | 127.4 | 0.63 | NA |
| 25%<br>Strength<br>TRIZol |  | 4.1 | 31.13 | 1.27 | 4.3 | 584.78 | 1.4 | 1.11 | 213.93 | 1 | 0.48 | 219.36 | 6.123 | 0.72 | 223 | 0.89 | NA |
|  |  | 4.2 | 103.58 | 1.2 | 0.59 | 235.8 | 1.03 | 0.46 | 256.84 | 0.99 | 0.45 | 288.25 | 6.421 | 0.9 | 149.1 | 0.64 | NA |
|  |  | 4.3 | 88.04 | 1.22 | 0.52 | 213.78 | 1.31 | 0.72 | 303.55 | 0.92 | 0.47 | 159.65 | 4.841 | 0.66 | 198.3 | 0.77 | NA |
| 0%<br>Water only |  | 5.1 | 44.4 | 1.35 | 1.31 | 21.73 | 1.78 | 3.29 | 175.37 | 1.15 | 0.48 | 6.47 | 0.648 | 0.2 | 109.3 | 1.46 | NA |
|  |  | 5.2 | 89.28 | 1.42 | 0.67 | 196.66 | 1.88 | 1.38 | 302.3 | 1.46 | 0.86 | 6 | 0.07 | 1.72 | 63 | 1.67 | NA |
|  |  | 5.3 | 139.83 | 1.28 | 1.43 | 53.23 | 2.43 | 9.14 | 87.06 | 1.5 | 0.68 | -2.83 | 0.161 | 0.35 | 110.3 | 41.58 | NA |

| Protein | Smpl | Francisella tularensis<br>subsp. holarctica LVS |  | Bacillus anthracis<br>Sterne strain |  | Yersinia enterocolitica |  | Mycobacterium<br>marinum |  | Burkholderia cepacia |  |
| --- | --- | --- | --- | --- | --- | --- | --- | --- | --- | --- | --- |
|  |  | Con.<br>mg/ml | 260/280 | Con.<br>mg/ml | 260/280 | Con.<br>mg/ml | 260/280 | Con.<br>mg/ml | 260/280 | Con.<br>mg/ml | 260/280 |
| 100% TRIzol | 1.1 | 0.13 | 1.17 | 6.3 | 1.48 | 2.13 | 1.47 | 3.73 | 1.46 | 0.48 | 1.44 |
|  | 1.2 | 0.46 | 1.28 | 4.07 | 1.54 | 1.07 | 1.55 | 0.95 | 1.48 | 0.39 | 1.53 |
|  | 1.3 | 0.39 | 1.32 | 3.63 | 1.66 | 1.6 | 1.51 | 2.91 | 1.51 | 0.55 | 1.44 |
| 75% Strength<br>TRIzol | 2.1 | 0.46 | 1.51 | 1.64 | 1.49 | 0.39 | 1.78 | 0.43 | 1.47 | 0.83 | 1.52 |
|  | 2.2 | 0.2 | 1.44 | 1.73 | 1.35 | 0.64 | 1.65 | 0.69 | 1.45 | 0.48 | 1.46 |
|  | 2.3 | 0.25 | 1.36 | 2.25 | 1.58 | 0.74 | 1.6 | 0.77 | 1.46 | 0.47 | 1.38 |
| 50% Strength<br>TRIzol | 3.1 | 0.86 | 1.64 | 2.57 | 1.56 | 0.98 | 0.85 | 0.97 | 1.39 | 0.41 | 1.31 |
|  | 3.2 | 0.91 | 1.39 | 3.68 | 1.52 | 0.82 | 1.08 | 0.11 | 1.42 | 0.56 | 1.18 |
|  | 3.3 | 0.95 | 1.81 | 2.36 | 1.51 | 0.61 | 1.56 | 1.55 | 1.09 | 0.44 | 1.33 |
| 25% Strength<br>TRIzol | 4.1 | 0.2 | 1.56 | 0.56 | 1.22 | 0.66 | 1.32 | 0.19 | 1.07 | 0.34 | 1.56 |
|  | 4.2 | 0.46 | 1.55 | 0.25 | 1.23 | 0.49 | 1.18 | 1.28 | 0.9 | 0.42 | 1.09 |
|  | 4.3 | 0.3 | 1.96 | 0.8 | 1.28 | 0.39 | 1.45 | 1.09 | 0.93 | 0.4 | 1.12 |
| 0%<br>Water only | 5.1 | 0.26 | 1.94 | 0.11 | 1.27 | 1.84 | 1.38 | 0.27 | 1.06 | 0.08 | 1.47 |
|  | 5.2 | 2.37 | 1.39 | 0.02 | 1.6 | 1.18 | 1.55 | 1.12 | 0.98 | 0.65 | 1.78 |
|  | 5.3 | 0.1 | 1.23 | 0.29 | 1.4 | 2.24 | 1.47 | 0.3 | 1.16 | 0.24 | 1.75 |

## Notes

### Competing Interest Statement

The authors have declared no competing interest.

